# Dynamic evolution of small signaling peptide compensation in plant stem cell control

**DOI:** 10.1101/2022.01.03.474791

**Authors:** Choon-Tak Kwon, Lingli Tang, Xingang Wang, Iacopo Gentile, Anat Hendelman, Gina Robitaille, Joyce Van Eck, Cao Xu, Zachary B. Lippman

**Affiliations:** School of Biological Sciences, Cold Spring Harbor Laboratory, Cold Spring Harbor, New York 11724, USA; Department of Horticultural Biotechnology, Kyung Hee University, Yongin 17104, Republic of Korea; State Key Laboratory of Plant Genomics, National Center for Plant Gene Research, CAS-JIC Centre of Excellence for Plant and Microbial Science (CEPAMS), Institute of Genetics and Developmental Biology, Chinese Academy of Sciences, Beijing, China; University of Chinese Academy of Sciences, Beijing, China; Boyce Thompson Institute, Ithaca, NY 14853, USA; Plant Breeding and Genetics Section, School of Integrative Plant Science, Cornell University, Ithaca, NY 14853, USA; Howard Hughes Medical Institute, Cold Spring Harbor Laboratory, Cold Spring Harbor, NY 11724, USA

**Keywords:** Paralogs, Redundancy, Epistasis, Signaling Peptide, *cis*-regulatory, Meristem, Tomato, Solanaceae, CRISPR

## Abstract

Gene duplications are a hallmark of plant genome evolution and a foundation for genetic interactions that shape phenotypic diversity^1–5^. Compensation is a major form of paralog interaction^6–8^, but how compensation relationships change as allelic variation accumulates is unknown. Here, we leveraged genomics and genome editing across the Solanaceae family to capture the evolution of compensating paralogs. Mutations in the stem cell regulator *CLV3* cause floral organs to overproliferate in many plants^9–11^. In tomato, this phenotype is partially suppressed by transcriptional upregulation of a closely related paralog^12^. Tobacco lost this paralog, resulting in no compensation and extreme *clv3* phenotypes. Strikingly, the paralogs of petunia and groundcherry nearly completely suppress *clv3*, indicating a potent ancestral state of compensation. Cross-species transgenic complementation analyses show this potent compensation partially degenerated in tomato due to a single amino acid change in the paralog and *cis*-regulatory variation that limits its transcriptional upregulation. Our findings show how genetic interactions are remodeled following duplications, and suggest that dynamic paralog evolution is widespread over short time scales and impacts phenotypic variation from natural and engineered mutations.

---

Gene duplications arise from whole genome and small-scale duplications and are pervasive in plant genomes^3,5,13,14^. Paralogs that emerge from duplications are completely redundant, which allows genetic variation to accumulate under relaxed selection^3,5^. This mutational drift can diversify paralog relationships through gene loss (pseudogenization), partitioning of ancestral functions (subfunctionalization), or gain of novel functions (neofunctionalization)^1,3,5,15^. Another prominent but less understood path of paralog evolution leads to “active compensation”, a form of redundancy where one or more paralogs are transcriptionally upregulated to substitute for the compromised activity of another^6,16,17^. Such relationships provide robustness against genetic or environmental change and may be under selection^18,19^. However, an often underappreciated paradox is that while duplications initially provide redundancy, they also promote new genetic variation through relaxed purifying selection^18,20,21^. Such variation, which can accumulate across both coding and *cis*-regulatory sequences, is the foundation for the broadly studied end-points of paralog diversification. What remains unclear is how such diversification modifies paralog functional relationships as species diversify over shorter time frames. This is because functional dissections of paralogs have been limited to within individual systems or between a few widely divergent species, and thus have failed to capture the trajectories and functional consequences of evolving compensatory relationships following lineage-specific ancestral duplications^6,12,14^.

*CLAVATA3/EMBRYO-SURROUNDING REGION-RELATED* (*CLE*) genes comprise an important gene family in plants encoding small-signaling peptides with diverse roles in growth and development^22,23^. CLE peptides are 12- or 13-residue glycopeptides processed from pre-propeptides^23,24^. The number and functional relationships, including redundancy, of CLE family members, vary considerably between distantly related species, due to lineage-specific duplications and variation in paralog retention and diversification^22^. However, the founding member from *Arabidopsis thaliana* (arabidopsis), CLAVATA3 (CLV3), is deeply conserved^9,25^. The CLV3 dodecapeptide is a ligand for the leucine-rich receptor kinase CLV1 and related receptors, and functions in a negative feedback circuit with WUSCHEL (WUS), a homeobox transcription factor that promotes stem cell production in shoot meristems^10,11^. Mutations in *CLV3* and its orthologs in many species cause meristem enlargement, which leads to tissue and organ overproliferation, or fasciation, phenotypes, especially in flowers^9,10^. We previously showed that *clv3* mutations in the divergent species arabidopsis, *Zea mays* (maize), and *Solanum lycopersicum* (tomato) are buffered through redundancy, but through different mechanisms^12^. In arabidopsis, multiple *CLE* family members partially suppress *clv3* without changing their expression^12^. In contrast to this “passive compensation”, a similar partial suppression of *clv3* mutations in maize (*zmcle7*) and tomato (*slclv3*) is achieved by active compensation from closely related *CLV3* paralogs^12^. Though the mechanism of compensation is shared between maize and tomato, the paralogs involved arose through lineage-specific duplications, indicating independent evolution of active compensation. Thus, it remains unclear how states of active compensation are achieved in any lineage and whether they remain stable or continue to evolve as species diversify.

With several genetically tractable species, closely related Solanaceae family members comprise a useful system to track the evolution of the compensation relationship between *CLV3* and its paralog. The compensating paralog in tomato, *SlCLE9*, originated from a duplication event just prior to diversification of the Solanales^12^. CRISPR-Cas9 engineered *slcle9* mutations result in normal plants, but strongly enhance *slclv3* due to loss of active compensation (**Fig. 1a-c**). Interestingly, our synteny analysis of 29 Solanaceae genomes capturing ~30 million years of evolution revealed several species that partially or completely lost their *SlCLE9* orthologs (**Fig. 1d and Supplementary Table 1**)^12^. For example, whereas *Physalis grisea* (groundcherry) and *Petunia hybrida* (petunia) have *SlCLE9* orthologs, *Capsicum annuum* (pepper) harbors only fragments of an *SlCLE9* ortholog, indicating pseudogenization (**Fig. 1d and Supplementary Table 1**)^12^. Both *S. tuberosum* (potato) and *S. melongena* (eggplant) lack *SlCLE9* orthologs entirely, and this presence-absence variation extends to the genus level; in *Nicotiana* (tobacco), the *SlCLE9* orthologs in *N. tabacum* and *N. benthamiana* were retained or pseudogenized, respectively (**Fig. 1d and Supplementary Table 1**).

**Fig. 1.**
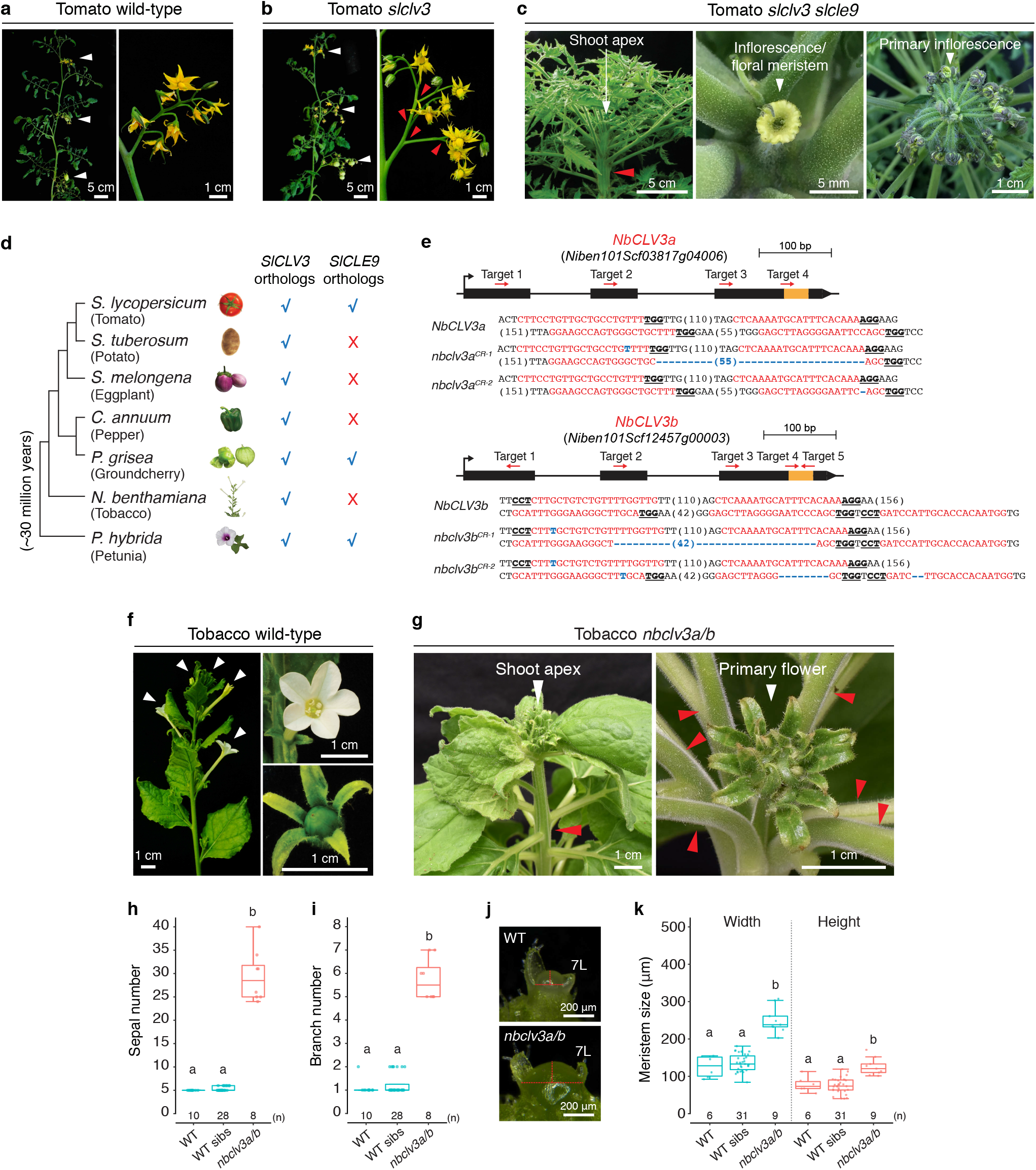
Loss of the tobacco *SlCLE9* ortholog abolished compensation. **a**, Shoot and inflorescence of tomato wild-type (WT). White arrowheads, inflorescences. **b**, Shoot and inflorescence of tomato *slclv3*. White arrowheads, inflorescences; red arrowheads, branches. **c**, Side and top-down view of tomato *slclv3 slcle9* shoot, inflorescence/floral meristem, and primary inflorescence. The red arrowhead indicates a fasciated shoot stem. **d**, Presence-absence variation of *SlCLE9* orthologs in the Solanaceae. Phylogenic relationships of the indicated species are shown. The blue checkmarks and the red Xs indicate presence and absence of the orthologs, respectively. **e**, Gene structures, and CRISPR-generated mutations of *NbCLV3a and NbCLV3b*. Orange rectangles indicate the CLE dodecapeptides regions. Targeted guide RNA (gRNA) and protospacer-adjacent motif (PAM) sequences are highlighted in red and bold underlined, respectively. Blue letters and dashes indicate insertions and deletions, respectively. Numbers in parentheses represent gap lengths. DNA sequences of gRNA target site 2 for both *NbCLV3a* and *NbCLV3b* are identical. **f**, Shoot, flower, and fruit pod of tobacco WT. White arrowheads, flowers. **g**, Side and top-down views of *nbclv3a/b* null mutants showing the shoot and primary flower. Red arrowheads indicate fasciated primary shoot (left panel) and shoot branches (right panel). **h**, Sepal number of primary flower from tobacco WT, WT sibling plants (WT sibs) and *nbclv3a/b* plants. **i**, Branch number of WT, WT sibs and *nbclv3a/b*. **j**, Primary shoot apical meristems from WT and *nbclv3a/b*. Red dotted lines mark width and height for meristem size quantification. 7L, 7^th^ leaf primordium. **k**, Quantification of meristem width and height from WT, WT sibs and *nbclv3a/b*. Box plots, 25^th^-75^th^ percentile; center line, median; whiskers, full data range in **h**, **i** and **k**. Exact sample sizes (n) for replicate types are indicated in **h**, **i** and **k**. Letters indicate significance groups at *P* < 0.01 (One-way ANOVA and Tukey test) in **h**, **i** and **k**. WT sibs are a mix of *nbclv3b* and *nbclv3a/+ nbclv3b* genotypes, which show wild-type phenotypes in **h**, **i** and **k**. See Source Data 4.

Since active compensation is typically mediated by the existence of a close paralog^6,16^, we predicted that species that lost their *SlCLE9* orthologs would lack active compensation. However, in such species, compensation could also have evolved from one or more *CLE* homologs, which could potentially compensate passively (i.e. without transcriptional upregulation), as found in the Brassicaceae species *Arabidopsis thaliana*^12^. We tested compensation in the allotetraploid *N. benthamiana*, where CRISPR-Cas9 genome editing is highly efficient, but brings an added layer of genetic complexity from having two sub-genome copies (orthologs) of all genes, including *NbCLV3* (*NbCLV3a* and *NbCLV3b*)^26^. To test for loss of compensation in this species, we designed a multiplex CRISPR-Cas9 construct with eight gRNAs designed to target *NbCLV3a* and *NbCLV3b* (four gRNAs each; **Fig 1e**). We obtained five first-generation transgenic (T_0_) plants, and unsurprisingly, all were chimeric (**Supplementary Fig. 1a-c**). Three of these plants exhibited severe fasciation phenotypes like tomato *slclv3 slcle9* double mutants, including thick stems and extreme overproliferation of floral organs, whereas the other two plants were less fasciated (**Supplementary Fig. 1c-d**). Though all plants were chimeric for mutations in *NbCLV3a* and *NbCLV3b*, sequencing showed the three strongest mutants carried only mutated alleles of both genes, suggesting a null-equivalent phenotype similar to tomato *slclv3 slcle9* double mutants (**Fig. 1c and Supplementary Fig. 1a-c**). To confirm that *NbCLV3a* and *NbCLV3b* function in a canonical *CLV-WUS* negative feedback circuit, we evaluated expression of both genes from shoot apices of individual T_0_ plants compared to wild-type (WT). Consistently, both genes were upregulated, similar to *SlCLV3* in *slclv3* mutants (**Supplementary Fig. 1e**)^12^. Though the severity of the floral fasciation in the strongest T_0_ plants precluded recovery of mutant seeds, these observations supported the absence of active compensation in *N. benthamiana*. Importantly, we further validated these results in T_1_ segregating lines derived from the weaker T_0_ plants, which fortuitously provided progeny populations that carried null alleles of *nbclv3b* and segregated for a null allele of *nbclv3a* (**Fig. 1e-i**). We used these populations to isolate *nbclv3a/b* allotetraploid mutants and showed that meristems were more than twice as large in these plants compared to *nbclv3b* single mutants and WT controls (**Fig. 1j, k**). Together, these results show that active compensation in the regulation of meristem maintenance was lost in *N. benthamiana* and also supports that conservation of active compensation in the Solanaceae requires retention of *SlCLE9* orthologs.

We next asked if compensation varies in lineages that retained their *SlCLE9* orthologs, and where allelic variation between these lineages could affect paralog function. Orthologous CLE pre-propeptide sequences are highly variable between species, but their dodecapeptides are more conserved^22,23^. Indeed, while *SlCLV3* and *SlCLE9* ortholog dodecapeptide sequences were nearly invariant in the Solanaceae, we found widespread variation in the coding and putative cis-regulatory regions of both genes, as determined by conserved non-coding sequence (CNS) analyses (**Supplementary Fig. 2 and Supplementary Table 1**). To assess active compensation in other Solanaceae species carrying *SlCLE9* orthologs, we took advantage of established CRISPR-Cas9 genome editing in petunia (**Fig. 2a**). Strikingly, the phenotypes of independently derived *phclv3* null mutants were both substantially weaker than tomato *slclv3* mutants (**Fig. 1b, 2b-d**). Although the primary shoot meristem was larger than WT meristems, 80% of *phclv3* flowers produced wild-type organ numbers (**Fig. 2c-f**). Given that multiple attempts to generate *pgcle9* mutants were unsuccessful, we micro-dissected *phclv3* meristems for RNA-sequencing to profile differentially expressed genes due to mutation of *PhCLV3*. Notably, out of all petunia *CLE* family members only *PhCLE9* was dramatically upregulated (>15-fold) (**Fig. 2g, h and Supplementary Table 2**), consistent with *SlCLE9* upregulation in tomato *slclv3* mutants and suggesting active compensation in petunia is mediated by *PhCLE9* and is stronger than in tomato.

**Fig. 2.**
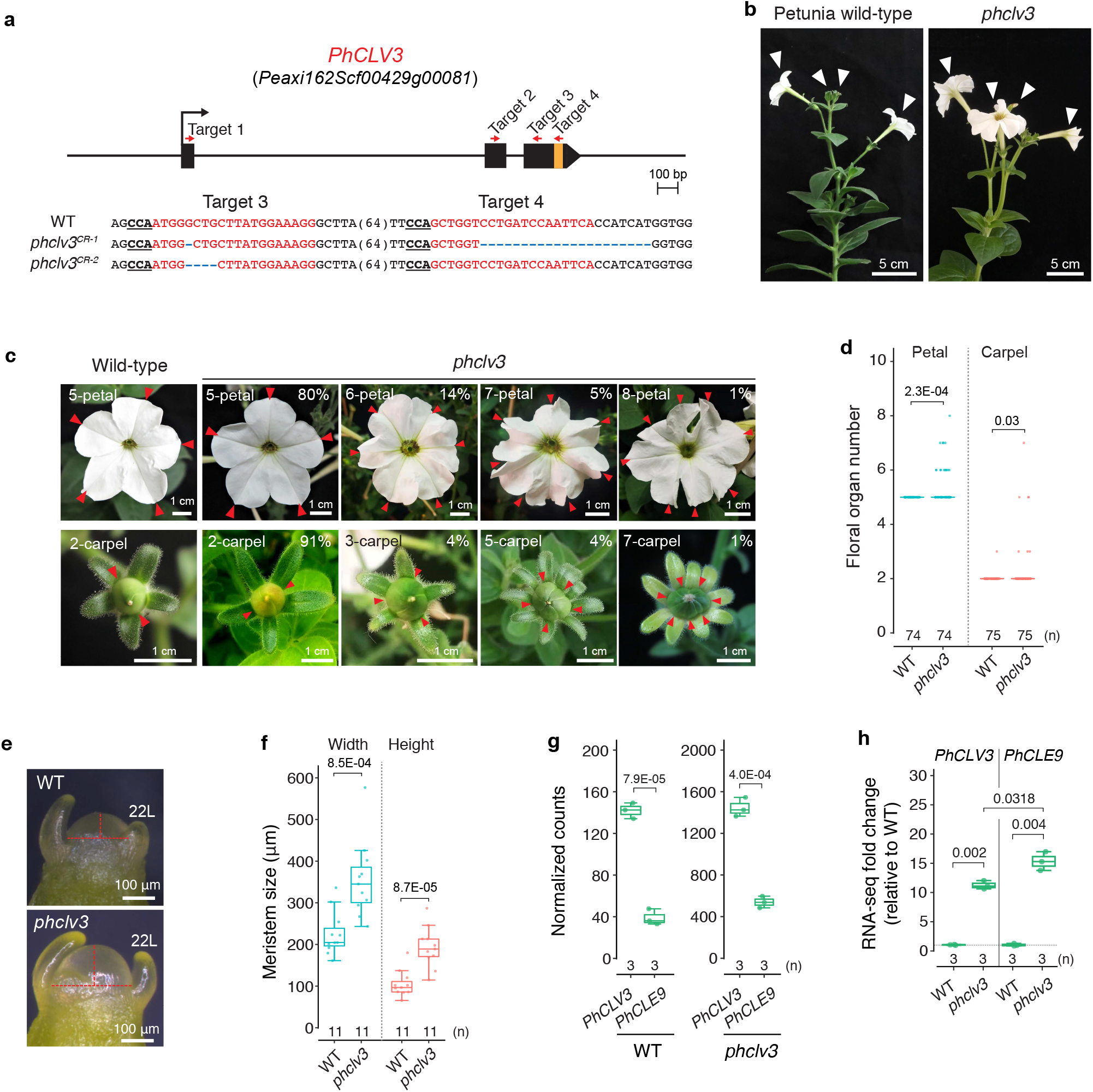
Weak fasciation of *phclv3* mutants in petunia indicates more potent compensation. **a**, Gene structure and sequences of two *phclv3* null alleles. Guide RNA and PAM sequences are highlighted in red and bold underlined, respectively. The orange rectangles in the gene structures represent the regions for CLE dodecapeptides. Numbers in parentheses represent gap lengths. Blue dashes indicate deletions. **b**, Shoot of petunia WT and *phclv3* plants. White arrowheads, flowers. **c**, Representative flowers and fruit pods of petunia WT and *phclv3* plants. Red arrowheads mark petals or carpels. Percentages indicate the proportions of flower and pod phenotypes. **d**, Quantification of petal and carpel numbers of WT and *phclv3*. **e**, Primary shoot apical meristems from petunia WT and *phclv3*. Red dotted lines mark width and height for meristem size quantification. 22L, 22^th^ leaf primordium. **f**, Quantification of meristem width and height from petunia WT and *phclv3*. **g**, Normalized read counts of *PhCLV3* and *PhCLE9* from WT and *phclv3* meristems. Expression unit, fragments per kilobase of transcript per million (FPKM). **h**, Expression fold-change of *PhCLV3* and *PhCLE9* relative to the normalized counts of WT from *phclv3*. Box plots, 25^th^-75^th^ percentile; center line, median; whiskers, full data range in **d**, **f**, **g** and **h**. *P* values (two-tailed, two-sample *t*-test) in **d**, **f**, **g** and **h**. Exact sample sizes (n) are shown as discrete numbers in **d**, **f**, **g** and **h**. Each replicate (n) is from 50-60 meristems in **g** and **h**.

Conservation of CLE dodecapeptide sequences is critical for proper ligand folding and receptor binding^27,28^. A single amino acid at position 6 distinguishes the petunia PhCLE9 and tomato SlCLE9 dodecapeptides, and a deeper analysis of conservation revealed that all species from tomato and its wild relatives through *Jaltomata sinuosa* have a serine at this position, whereas all other Solanaceae except for a subset of tobacco species have a glycine (**Fig. 3a, Supplementary Fig. 2c and Supplementary Table 1**)^12,22^. Beyond the Solanaceae, this glycine is invariant in angiosperm CLV3 orthologs, is highly conserved in other CLE peptides, and is essential in Arabidopsis CLV3 and CLE41 peptides for precise binding to their receptors (**Supplementary Fig. 2 and Supplementary Table 1**)^12,22,27–30^. These observations suggested that other Solanaceae species with the conserved glycine in their *SlCLE9* orthologs might have more effective ligands, and would also be more potent compensators than tomato *SlCLE9*. We tested this using CRISPR-Cas9 genome editing in groundcherry (**Supplementary Fig. 3**). Notably, null mutation of groundcherry *pgclv3* resulted in only weak phenotypes similar to petunia *phclv3* mutants (Fig 3b-e and Supplementary Fig. 3a, b). We also engineered homozygous *pgcle9* null mutations, which were nearly identical to wild-type (**Fig. 3b-e and Supplementary Fig. 3c**), and consistent with these weak effects, the sizes of primary shoot meristems in both mutants were largely unchanged (**Fig. 3f, g**). Importantly, as in tomato and in petunia, the expressions of both *PgCLV3* and *PgCLE9* were upregulated in *pgclv3* meristems (**Fig. 3h, i and Supplementary Table 3**), and *pgclv3 pgcle9* double null mutants were severely fasciated, similar to tomato *slclv3 slcle9* double mutants, confirming conservation of active compensation (**Fig. 3j, k and Supplementary Fig. 3d, e**). Thus, while active compensation is conserved between tomato, petunia, and groundcherry, compensation from *SlCLE9* orthologs in petunia and groundcherry is stronger than in tomato.

**Fig. 3.**
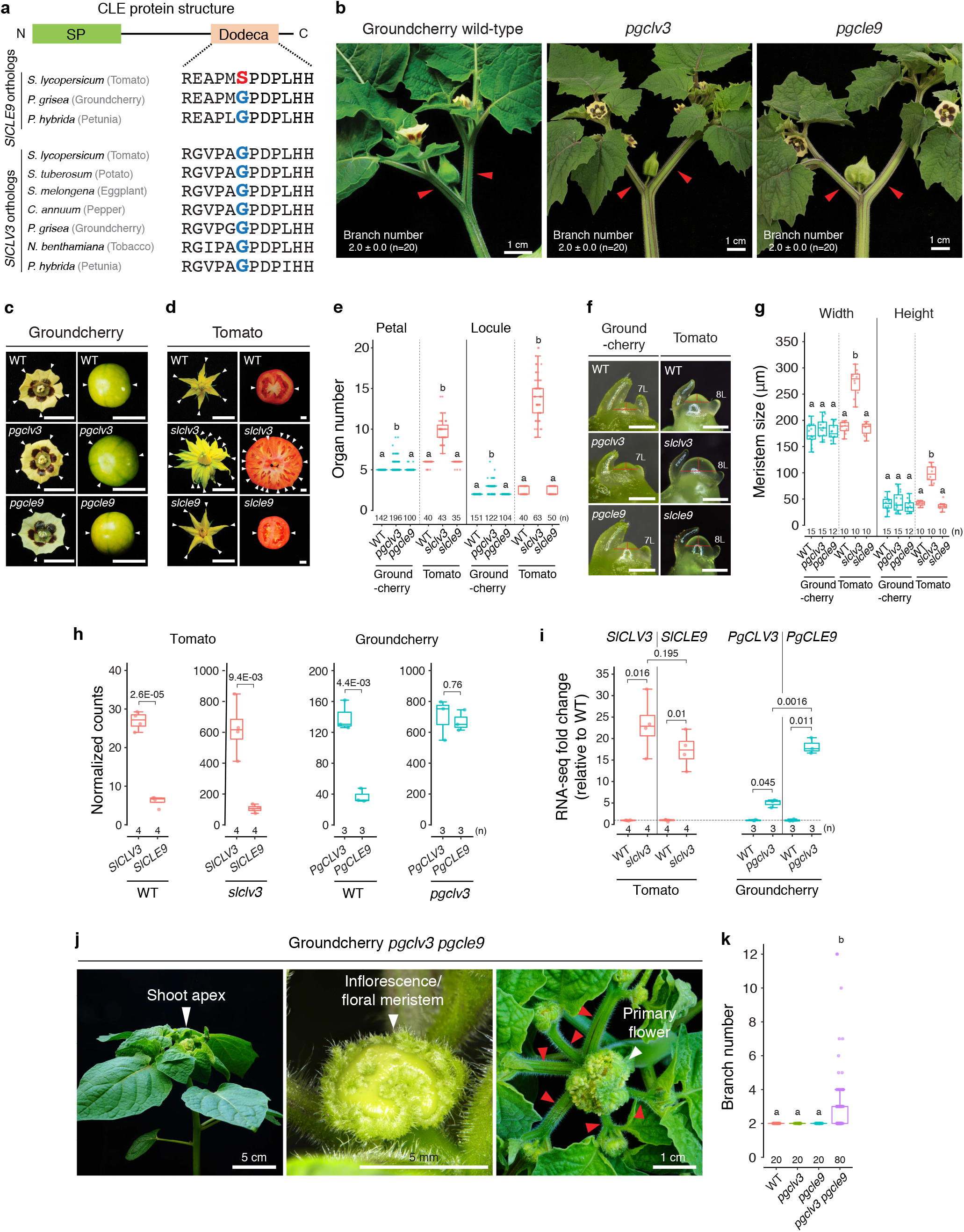
A highly conserved dodecapeptide amino acid is associated with potent compensation in groundcherry. **a**, CLE protein structure and dodecapeptide sequences of *SlCLE9* and *SlCLV3* orthologs in the Solanaceae. The sixth residue of the dodecapeptides are highlighted by red or blue bold font. **b**, Shoot and inflorescences of groundcherry WT, *pgclv3* and *pgcle9* plants. Red arrowheads mark two side shoots that develop after single-flowered inflorescences. **c**, Representative flowers and fruits from groundcherry WT, *pgclv3*, and *pgcle9* plants. Scale bar, 1 cm. **d**, Representative flowers and fruits from tomato WT, *slclv3*, and *slcle9* plants. White arrowheads mark petals or locules. Scale bar, 1 cm. **e**, Quantification and comparison of petal and locule numbers from groundcherry WT, *pgclv3, pgcle9* and tomato WT, *slclv3*, and *slcle9* plants. **f**, Primary shoot apical meristems from groundcherry WT, *pgclv3, pgcle9* and tomato WT, *slclv3*, and *slcle9* plants. 7L, 8L: 7^th^ and 8^th^ leaf primordia, respectively. Red dotted lines indicate width and height for meristem size measurements, Scale bar, 200 μm. **g**, Quantification of meristem width and height from groundcherry WT, *pgclv3, pgcle9*, tomato WT, *slclv3*, and *slcle9* plants. **h**, Normalized RNA-seq read counts of *SlCLV3*, *SlCLE9*, *PgCLV3*, and *PgCLE9* from tomato WT, *slclv3*, groundcherry WT and *pgclv3* meristems. Expression units are counts per million (CPM) for tomato and transcripts per million (TPM) for groundcherry **i**, Expression fold-change of *SlCLV3*, *SlCLE9*, *PgCLV3*, and *PgCLE9* relative to the normalized counts of WT expressions of these genes in the indicated genotypes. **j**, Side and top-down views of a *pgclv3 pgcle9* double mutant shoot, inflorescence/floral meristem, and primary flower. Red arrowheads indicate branches that emerged after the primary flower. **k**, Branch number of WT, *pgclv3, pgcle9*, and *pgclv3 pgcle9* plants. Box plots, 25^th^-75^th^ percentile; center line, median; whiskers, full data range in **e**, **g**, **h**, **i** and **k**. The letters indicate the significance groups at *P* < 0.01 (One-way ANOVA and Tukey test) in **e**, **g** and **k**. *P* values (two-tailed, two-sample *t*-test) in **h** and **i**. Exact sample sizes (n) are shown in **e**, **g**, **h**, **i** and **k**. Each replicate (n) is from 30-35 meristems in **h** and **i**.

Our dissections of active compensation in tomato, petunia, and groundcherry suggested that the conserved glycine of the dodecapeptide is necessary for potent compensation. In further support, two conserved residues (Aspartic acid and Phenylalanine) in SlCLV1, which is the primary receptor of SlCLV3 and SlCLE9 ligands^12^, are critical for interaction with the sixth glycine of CLE peptides (**Supplementary Fig. 4**)^29,30^. Solanaceae CLV1 orthologs are invariant in these ligand binding residues (**Supplementary Fig. 4**). To test if the groundcherry and petunia orthologs of CLV1 (PgCLV1 and PhCLV1) are also the primary receptors for PgCLE9 and PhCLE9 as in tomato, we made double mutants between the weakly fasciated groundcherry *pgclv1* and *pgclv3* and also the weakly fasciated petunia *phclv1* and *phclv3* null mutants (**Supplementary Fig. 5)**^31^. Consistently, the double null mutants in both species matched the severe fasciation of groundcherry *pgclv3 pgcle9* double mutants, and importantly, also the tomato *slclv1 slclv3* and *slclv3 slcle9* double mutants (**Fig. 1c, 3j and Supplementary Fig. 5c-e**). These results support that the glycine to serine change in the tomato SlCLE9 dodecapeptide could be reducing binding affinity to SlCLV1, thus explaining weaker compensation in this species.

To test the significance of the glycine, we asked if the genomic sequence of *PgCLE9* (*gPgCLE9^PgCLE9^*) could complement *slclv3* mutants (**Fig. 4a**). While *slclv3* fasciation is nearly completely suppressed by the genomic sequence of *SlCLV3* (*gSlCLV3^SlCLV3^*), *gPgCLE9^PgCLE9^* had no effect (**Fig. 4a, b and Supplementary Fig. 6a, b**). Poor heterologous expression between groundcherry and tomato could explain this result, so we transformed *slclv3* mutants with a construct expressing the groundcherry dodecapeptide from the genomic sequence of tomato *SlCLE9* (*gSlCLE9^PgCLE9^*) (**Fig. 4a, b and Supplementary Fig. 6a, b**). Surprisingly, this construct also failed to complement, leading us to ask if strong active compensation depended on the conserved glycine as well as higher expression of dodecapeptides having the glycine. In support of this, in contrast to tomato, the fold-change increases in expression of both groundcherry *PgCLE9* and petunia *PhCLE9* were higher relative to upregulation of *CLV3* in their respective *clv3* mutants (**Fig. 2h, 3i**). As the promoter of tomato *SlCLV3* is more transcriptionally responsive than the promoter of *SlCLE9* to *slclv3* mutations (**Fig. 3h**), we used a construct expressing the groundcherry dodecapeptide from *SlCLV3* genomic sequence (*gSlCLV3^PgCLE9^*), which strongly suppressed *slclv3* mutants. Notably, this complementation was slightly weaker than with *gSlCLV3^SlCLV3^*, consistent with active compensation from PgCLE9 and PhCLE9 dodecapeptides in groundcherry and petunia still permitting weak phenotypes of their respective *clv3* mutants (**Fig. 4a, b and Supplementary Fig. 6a, b**). A construct expressing the tomato SlCLE9 dodecapeptide from the same *SlCLV3* genomic sequence (*gSlCLV3^SlCLE9^*) failed to complement, indicating that higher expression alone is insufficient (**Fig. 4a, b and Supplementary Fig. 6a, b**). Consistently, a weaker expression of PgCLE9 dodecapeptide (*gSlCLE9^SlCLE9S6G^*) or a stronger expression of SlCLE9 dodecapeptide (*gSlCLV3^SlCLE9^-2*) could only suppress *slclv3 slcle9* double mutants to *slclv3* single mutant phenotypes (**Supplementary Fig. 6c, d)**. Altogether, our results show that changes in both the dodecapeptide and its expression explain evolutionary variation in the strength of compensation between tomato and its relatives groundcherry and petunia (**Fig. 4c**).

**Fig. 4.**
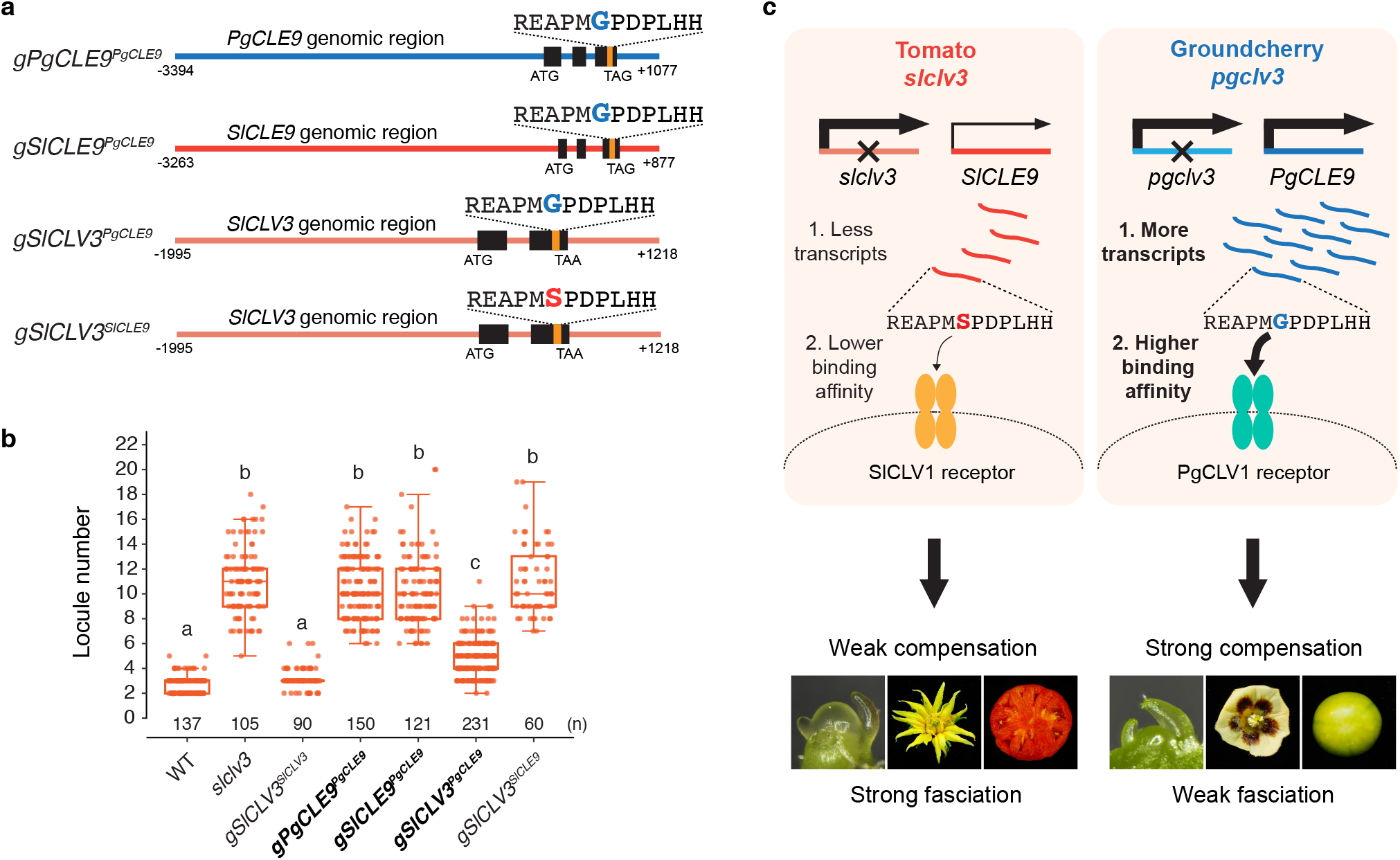
Variation in Solanaceae compensation is due to changes in both the SlCLE9 ortholog dodecapeptide and its expression. **a**, Diagrams of constructs used for complementation tests. *gPgCLE9^PgCLE9^* (*PgCLE9* genomic DNA). *gSlCLE9^PgCLE9^* (*SlCLE9* genomic DNA including the sequence for PgCLE9 dodecapeptide). *gSlCLV3^PgCLE9^* (*SlCLV3* genomic DNA including the sequence for PgCLE9 dodecapeptide). *gSlCLV3^SlCLE9^* (*SlCLV3* genomic DNA including the sequence for SlCLE9 dodecapeptide). Black and orange rectangles mark the coding sequences and the dodecapeptide sequences, respectively. The numbers with minus (-) and plus (+) signs indicate the positions of the upstream sequences and the downstream sequences from the adenines of start codons, respectively. **b**, Locule number quantification from WT and *slclv3* mutants compared to T_1_ transgenic plants of *gSlCLV3^SlCLV3^*, *gPgCLE9^PgCLE9^, gSlCLE9^PgCLE9^, gSlCLV3^PgCLE9^*, and *gSlCLV3^SlCLE9^*. Box plots, 25^th^-75^th^ percentile; center line, median; whiskers, full data range. The letters indicate the significance groups at *P* < 0.01 (One-way ANOVA and Tukey test). Exact sample sizes (n) are shown as discrete numbers. Data are based on at least 10 independent transgenic lines for each construct. **c**, A proposed model for differences in active compensation between tomato and groundcherry. The more potent active compensation in groundcherry compared to tomato is due to both the glycine-containing PgCLE9 dodecapeptide and its higher expression.

Here, we uncovered a dynamic evolution of paralogs interacting in an active compensation relationship. A first step of paralog diversification that can promote their preservation is ‘compensatory drift’, through which optimal levels of dosage-sensitive genes are maintained by reducing the expression of one paralog and elevating the other^32^. *CLV3* orthologs are dosage-sensitive^33–35^, and the consistently higher expression levels of Solanaceae *CLV3* orthologs relative to *SlCLE9* orthologs indicate that compensatory drift and active compensation emerged soon after duplication (**Fig. 2g, 3h**). However, despite this expression rebalancing, we found that *CLV3* compensation degraded multiple times during the Solanaceae family radiation over the last ~30 million years (**Fig. 5**). At one extreme, *N. benthamiana*, and likely other species that lost their *SlCLE9* orthologs, completely lost active compensation and thus buffering of meristem homeostasis. In tomato, both coding and *cis*-regulatory changes weakened *SlCLE9*, and we pinpointed a critical amino acid change that facilitated partial degradation of compensation from the more potent ancestral state found in groundcherry and petunia (**Fig. 5**). Thus, the differential accumulation of genetic variation between *SlCLE9* orthologs in these four Solanaceae species resulted in both qualitative and quantitative differences in compensation potencies. Our finding of extensive coding and *cis*-regulatory variation between *SlCLE9* orthologs suggests a range of potencies could exist in Solanaceae *CLV3* compensation (**Supplementary Fig. 2 and Supplementary Table 1**). For example, even among tobacco species, while *N. benthamiana* lost compensation, *N. obtusifolia* likely has strong compensation due to retention of a glycine-containing *SlCLE9* ortholog, and surprisingly, the sub-genome copies of *SlCLE9* orthologs in *N. attenuata*, *N. tabacum*, and *N. tomentosiformis* each have a glycine and a serine (**Supplementary Fig. 2c and Supplementary Table 1**).

**Fig. 5.**
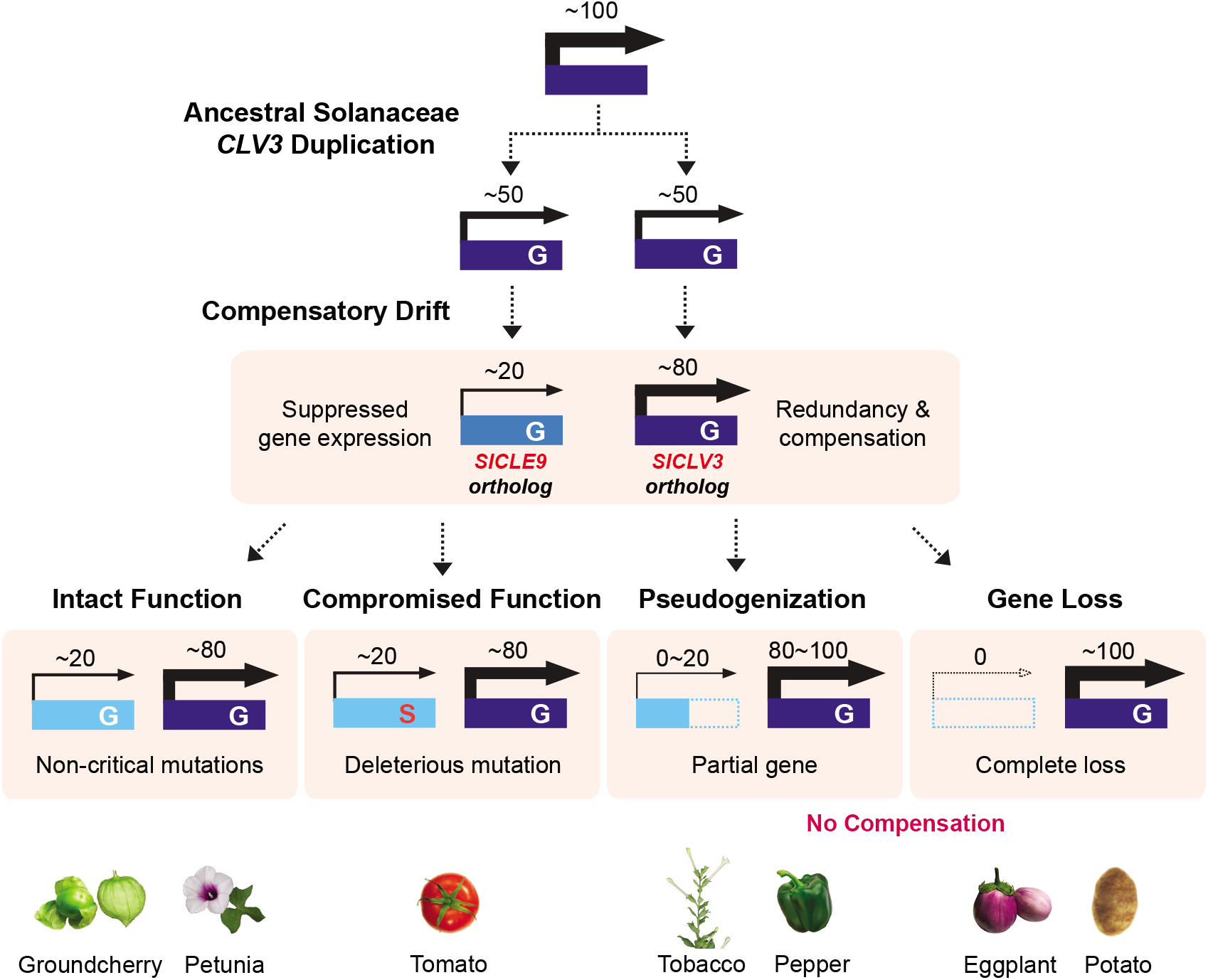
Summary and model of the dynamic evolution of *SlCLV3* and *SlCLE9* orthologs and their compensation relationships in the Solanaceae. Dark blue, blue, and sky blue rectangles indicate the coding region of the genes. Arrows and their thickness represent gene expressions and their relative levels, respectively. Numbers above the arrows indicate hypothetical relative proportions of *SlCLV3* and *SlCLE9* ortholog expression levels. ‘G’ and ‘S’ within the rectangles denote the sixth amino acid of each CLE dodecapeptide. Dashed rectangles mark deletions of the coding region, resulting in pseudogenes (pepper and tobacco) and complete gene loss (eggplant, potato) in each genome. The red gradient bar reflects the loss of active compensation and its degree, depending on the indicated genetic variation.

Differences in transcriptional control may play a larger role. Widespread variation in *cis*-regulatory regions among tomato species suggests even greater variation between species in the Solanaceae family^36^. Such diversity, both within and between genera (**Supplementary Fig. 2a**), could result in differences in upregulation of *SlCLE9* orthologs and phenotypes when *CLV3* activity is compromised. Such a wide range of compensation strengths could be a foundation for species-specific phenotypes. Notably, a structural variant that partially disrupts the promoter of *SlCLV3* is a major tomato domestication fruit size QTL, and we found that its severity was mitigated by active compensation from *SlCLE9*, resulting in a more moderate effect that may have facilitated selection^12,37^. The increase in fruit size from this variant may not have emerged if the ancestral version of *SlCLE9* was retained in tomato, and moreover, in groundcherry and other Solanaceae orphan crops with potent *SlCLE9* orthologs, engineering mutations in *CLV3* alone would likely not benefit fruit size^31,38^. Beyond the Solanaceae, variation in meristem shape and form is associated with morphological variation within and between species^39–41^. Such differences could in part be based on variation in compensation between meristem homeostasis genes, which could also influence phenotypic outcomes from engineered variation in CLV network genes^33,35,42^.

More broadly, our findings have important implications in understanding and exploiting phenotypic changes caused by natural and engineered variation in other species and gene families. The era of pan-genomes^43–46^ continues to uncover remarkable diversity in paralogs, including presence-absence variation, as well as widespread coding and regulatory variation between retained paralogs. Our findings show that such variation, much of which could be cryptic^47–49^, can impact phenotypes in unpredictable and subtle ways when members of a gene family are mutated within or between species. Revealing and dissecting diverse paralogous relationships can advance our understanding of how dynamically evolving duplicated genes shape phenotypic variation across short time scales, and improve predictability in trait engineering of both old and new crops.

## Methods

### Plant materials and growth conditions

Seeds of petunia (*P. hybrida* ‘W115’, Mitchel diploid) were provided by Prof. Yulong Guo, Southwest University (Chongqing, China). Seeds of tobacco (*N. benthamiana*), groundcherry (*P. grisea*) and tomato (*S. lycopersicum*, cultivar M82) were from Cold Spring Harbor Laboratory (CSHL) seed stocks. All seeds were sown directly in soil and grown in growth chambers, greenhouses or fields at CSHL, New York, USA (tomato, tobacco, groundcherry) and Institute of Genetics and Developmental Biology, Chinese Academy of Sciences, Beijing, China (petunia). Briefly, groundcherry and tomato seedlings were grown in the greenhouse or field at CSHL as described previously^50^. Tobacco plants were grown under long-day conditions (16 h light, 21°C/8 h dark, 20°C; 40-55% relative humidity; 75 μmol m^−2^ s^−1^) in the greenhouse at CSHL. *Petunia* plants were grown under long-day conditions (16h light, 25°C/8h dark, 21°C; 50-60% relative humidity; 75 μmol m^−2^ s^−1^) in growth chambers and greenhouses at Institute of Genetics and Developmental Biology, Chinese Academy of Sciences. All plants were grown under overhead watering (tobacco) or drip irrigation (groundcherry, petunia and tomato), and standard fertilizer regimes.

### CRISPR–Cas9 genome editing and plant transformation

Targeted mutagenesis using the CRISPR-Cas9 system for tobacco, groundcherry, and petunia were performed as described previously^31,51–57^. Briefly, the binary vectors were constructed through Golden Gate cloning as described^51,58^, and introduced into tobacco, groundcherry, and petunia by *Agrobacterium tumefaciens*-mediated transformation as described^52,53,57,59^. First-generation transgenic plants were transplanted in soil and genotyped to validate CRISPR-generated mutations by PCR and Sanger sequencing, as previously described^37^. All primer and gRNA sequences are included in **Supplementary Table 4**.

### Plant phenotyping and meristem imaging

All phenotypic quantification data on inflorescences and fruits were performed as previously described^12,37^. Briefly, the phenotypic characterization was performed with biallelic or chimeric T_0_ plants (tobacco), and non-transgenic homozygous plants (groundcherry, petunia, and tomato) from backcrossing or selfing. CRISPR-generated null mutants of groundcherry and tomato were sprayed with 400 mgl^−1^ kanamycin, and petunia were sprayed with 100mgl^−1^ kanamycin and genotyped by PCR to verify the absence of the transgenes. We manually counted the floral organs (petal and carpel/locule) from multiple inflorescence and plants. All the exact sample numbers of individual transgenic plants and aggregated organ quantifications are marked in the figures and are collated in the Source data. Meristem imaging and size quantification were conducted as described previously^37,60^. Briefly, the images of hand-dissected meristems were captured on a Nikon SMZ1500 (tomato), Nikon SMZ25 (groundcherry and tomato). Dissection and stereomicroscope imaging of petunia meristems were carried out under Olympus microscope (SteREO Discovery, v.12).

### RNA extraction, complementary DNA synthesis and quantitative real-time PCR (qPCR)

RNA extraction and qPCR for tobacco, groundcherry and petunia were conducted as previously described with minor modification^12,50^. Briefly, for total RNA of the tobacco apices, the dissected shoot apices were extracted with RNeasy Plant Mini Kit (QIAGEN) according to the manufacturer’s instructions. 1 μg total RNA was used for reverse transcriptase PCR with the SuperScript III First-Strand Synthesis System (Invitrogen). qPCR was conducted in the iQ SYBR Green Supermix (Bio-Rad Laboratories) reaction system on the CFX96 Real-Time PCR Detection System (Bio-Rad) with gene-specific primers (**Supplementary Table 4**)^61^. For total RNA of the groundcherry meristems, the hand-dissected shoot apical meristems were extracted by the ARCTURUS PicoPure RNA Extraction Kit (Applied Biosystems). 30–35 meristems were collected for one replicate for each genotype. Total RNA of the petunia meristems was also extracted by the ARCTURUS PicoPure RNA Extraction Kit (Applied Biosystems). 50–60 meristems were collected for one replicate for each genotype.

### Meristem transcriptome profiling

The transcriptome data from tomato meristems were obtained from our previous RNA-seq data deposited in the Sequence Read Archive project (SRP161864) and BioProject (PRJNA491365)^12^. RNA-seq and differentially expressed genes (DEGs) analyses of groundcherry and petunia meristems were performed as previously described with slight modification^12^. Briefly, the libraries for RNA-sequencing (RNA-seq) were prepared by the KAPA mRNA HyperPrep Kit (Roche). The quality of each library was validated with a 2100 Bioanalyzer (Agilent Technologies). Paired-end 75-base sequencing was conducted on the Illumina sequencing platform (NextSeq, Mid-Output). Reads for the wild-type (WT) groundcherry and *pgclv3* mutant were trimmed by quality using Trimmomatic v.0.32 (parameters: ILLUMINACLIP:TruSeq3-PE-2.fa:2:40:15:1:FALSE LEADING:30 TRAILING:30 MINLEN:50)^62^ and aligned to the reference transcriptome assembly of groundcherry^31^ for quantification using ‘kallisto quant’ (v0.46.2, bootstrap: 100)^63^. Kallisto quantification results were used as inputs for ‘sleuth’ in R to get normalized estimated counts for each transcript^64^. For RNA-seq of petunia meristems, the libraries were prepared by SMARTer Ultra Low Input RNA for Sequencing Kit (Clontech). The quality of each library was validated with a 2100 Bioanalyzer (Agilent Technologies). Paired-end 150-base sequencing was conducted on the Illumina NovaSeq 6000 sequencing platform (NextSeq, Mid-Output). Reads for the WT petunia and *phclv3* mutant were trimmed by quality using Trimmomatic v0.36 (ILLUMINACLIP:adapter.fa:2:30:10 LEADING:20 TRAILING:20 SLIDINGWINDOW:4:15 MINLEN:36)^62^ and aligned to the reference genome sequence of petunia^65^ using hisat2 v2.1.0 with default parameters^66^. Alignments were sorted with samtools v1.8^67^ and the RNA-seq reads were assembled using StringTie (v2.0.3) with default parameters^68^. To verify and annotate the transcript of petunia *PhCLE9* (Peaxi162Scf00429:766800-783916), orthologous Blast was performed using tomato *SlCLE9* as a bait and the resulting transcript was confirmed by PCR amplification followed by Sanger sequencing (see **Supplementary Dataset 2**). The expected read counts and fragments per kilobase of transcript per million mapped reads (FPKM) were also calculated using SringTie (v2.0.3)^68^. The statistical analyses for groundcherry and petunia data were performed in R(v3.5.2) (RStudio (v.1.1.463)) and R(v4.0.3), respectively^69,70^. Significant differential expression between groundcherry WT and *pgclv3* mutant was identified with sleuth^64^ using *q*-value ≤ 0.01 cut-offs. Significant differential expression between petunia WT and *phclv3* mutant was confirmed with DESeq2^64,71^ using *q*-value ≤ 0.05 and |log2_ratio| ≥ 1.

### Transgenic complementation of *PgCLE9, SlCLV3* and *SlCLE9*

The transgenic lines and genomic DNA sequence for *gSlCLV3^SlCLV3^* and *gSlCLV3^SlCLE9^* were procured from our previous study^12^. The genomic DNA sequences of *PgCLE9* consisted of *gPgCLE9^PgCLE9^* 4471 base pair (bp) in total with 3394 bp upstream, 548 bp of coding sequence containing introns, and 529 bp downstream. The genomic DNA sequences of *SlCLE9* consisted of *gSlCLE9^SlCLE9^* 4140 bp in total with 3263 bp upstream, 403 bp of coding sequence containing introns, and 474 bp downstream. Site-directed mutageneses were performed to substitute the SlCLE9 dodecapeptide into PgCLE9 within *gSlCLE9^SlCLE9^* (*gSlCLE9^PgCLE9^*) and the SlCLV3 dodecapeptide into PgCLE9 within *gSlCLV3^SlCLV3^* (*gSlCLV3^PgCLE9^*). The PCR products were amplified from the vectors including the genomic region of *SlCLV3* (pICH47742-*gSlCLV3^SlCLV3^*) and *SlCLE9* (pICH47742-*gSlCLE9^SlCLE9^-2*) with overlapping primers (**Supplementary Table 4**) using KOD One™ PCR Master Mix (TOYOBO). Then, the amplified PCR products were digested using DpnI (New England Biolabs) and transformed into DH5a competent cells. The sequences of the resulting plasmids were confirmed by Sanger sequencing with multiple primers (**Supplementary Table 4**). The Level 1 vectors (pICH47742-*gPgCLE9^PgCLE9^*, *gSlCLE9^PgCLE9^* and *gSlCLV3^PgCLE9^*) were assembled with the construct pICH47732-NOSpro::NPTII into the binary vector pICSL4723 through Golden Gate cloning as previously described^51,58,72^. The binary vectors were introduced into the tomato *slclv3* mutant by *Agrobacterium tumefaciens*-mediated transformation as previously described^53^. The genomic DNA sequences of *SlCLV3* consisted of *gSlCLV3^SlCLV3^-2* 3213 bp in total with 1995 bp upstream, 600 bp of coding sequence containing introns, and 618 bp downstream. The genomic DNA sequences of *SlCLE9* consisted of *gSlCLE9^SlCLE9^-2* 2740 bp in total with 1996 bp upstream, 403 bp of coding sequence containing introns, and 341 bp downstream. Site-directed mutagenesis was performed to substitute the SlCLV3 dodecapeptide into SlCLE9 within *gSlCLV3^SlCLV3^* (*gSlCLV3^SlCLE9^-2*) and the SlCLE9 dodecapeptide into SlCLE9^S6G^ within *gSlCLE9^SlCLE9^* (*gSlCLE9^SlCLE9S6G^*). The PCR products were amplified from the vectors including the genomic region of *SlCLV3* (pDONOR221-*gSlCLV3^SlCLV3^-2*) and *SlCLE9* (pDONOR221-*gSlCLE9^SlCLE9^* with overlapping primers (**Supplementary Table 4**) using KOD One™ PCR Master Mix (TOYOBO). Then, the amplified PCR products were digested using DpnI (New England Biolabs) and transformed into DH5a competent cells. The sequences of the resulting plasmids were confirmed by Sanger sequencing with multiple primers (**Supplementary Table 4**), and colonies were recombined into binary vector pGWB401^73^ for transgenic complementation. The binary vectors were introduced into the tomato *slclv3 slcle9* double mutant by *Agrobacterium tumefaciens*-mediated transformation as previously described^53^. Transgenic lines were confirmed by PCR and kanamycin resistance, and at least three independent transgenic lines from each construct were used for data collection (see **Source Data**).

### Conserved noncoding sequence (CNS) analysis

Analysis of conserved non-coding sequences (CNSs) is a common approach to identify putative *cis*-regulatory sequences of genes (e.g. promoters, enhancers). Solanaceae orthologous genes of *SlCLV3* and *SlCLE9* for synteny analysis and CNSs in the promoter regions surrounding the orthologs of *SlCLV3* and *SlCLE9* were identified using our previously developed Conservatory algorithm, using default parameters^74^. In parallel, all of the genomes were scanned with tBLASTn to find mis- or unannotated protein coding regions for each gene. CNSs in the promoter regions were called by Conservatory using default parameters^74^. To calculate protein identity percentages and dodecapeptide identity percentages, protein sequences were aligned by MAFFT (v.7.45) using BLOSUM62 matrix and ‘E-INS-i’ and ‘G-INS-i’ algorithm respectively^75^.

### Statistical analyses

Statistical calculations were conducted using R(v3.5.2 and v4.0.3)^69^ and Microsoft Excel, as previously described^50^. Statistical analyses were performed using a two-tailed, two-sample t-test and a one-way analysis of variance (ANOVA) with Tukey test. The exact sample sizes (n) and all raw data for each experimental group/condition are given as discrete numbers in each figure panel and Source data. Additional information is available in the Nature Research Reporting Summary, which includes statements on statistics, software used and data availability.

## Data availability

Raw data and information for CRISPR-generated alleles, all quantifications, synteny analysis, and exact *P* values (One-way ANOVA and Tukey test) are in **Source data**. The raw Sanger sequence traces for edited sequences are in **Supplementary Dataset 1**. The groundcherry and petunia BioProject accession numbers are PRJNA704671 and PRJNA750419, respectively.

## Acknowledgements

We thank members of the Lippman laboratory for comments and discussions, and also critical friends Y. Eshed and M. Bartlett. We thank S. Soyk for discussions and initial peptide sequence analysis in tomato, and J. Kim and A. Krainer for technical support. We thank A. Horowitz Doyle, K. Swartwood, M. Tjahjadi, L. Randall, and P. Keen from the Van Eck lab for performing tobacco, groundcherry and tomato transformations. We thank T. Mulligan, K. Schlecht, A. Krainer and S. Qiao for assistance with plant care. This research was supported by National Natural Science Foundation of China (grant no. 31972423 and 31991183) and Chinese Academy of Sciences (grant no. 153E11KYSB20180019) to C.X., the Howard Hughes Medical Institute, an Agriculture and Food Research Initiative competitive grant from the USDA National Institute of Food and Agriculture (grant no. 2016-67013-24452) and the National Science Foundation Plant Genome Research Program (grant no. IOS-1732253 and IOS-1546837) to Z.B.L.

## Author contributions

C.-T.K. designed the research and conducted the experiments, prepared the figures and wrote the manuscript. L.T. performed the petunia CRISPR experiments, tomato transgenic complementation tests, genetic, RNA-seq and phenotypic analyses. X.W. performed the groundcherry RNA-seq, phenotypic analyses, and wrote the manuscript. I.G performed the tobacco genetic and phenotypic analyses. A.H. performed CLE family analyses. G.R. characterized CRISPR mutations. J.V.E generated transgenic plants and CRISPR lines. C.X. supervised and led the petunia CRISPR experiments and tomato transgenic complementation tests, genetic, RNA-seq and phenotypic analyses, contributed ideas and edited the manuscript. Z.B.L. conceived and led the research, supervised and performed the experiments, prepared the figures and wrote the manuscript. All authors read, edited, and approved the manuscript.

## Competing interests

The authors declare that they have no competing interests.

## Additional information

### Supplementary figure legends

**Supplementary Fig. 1.**
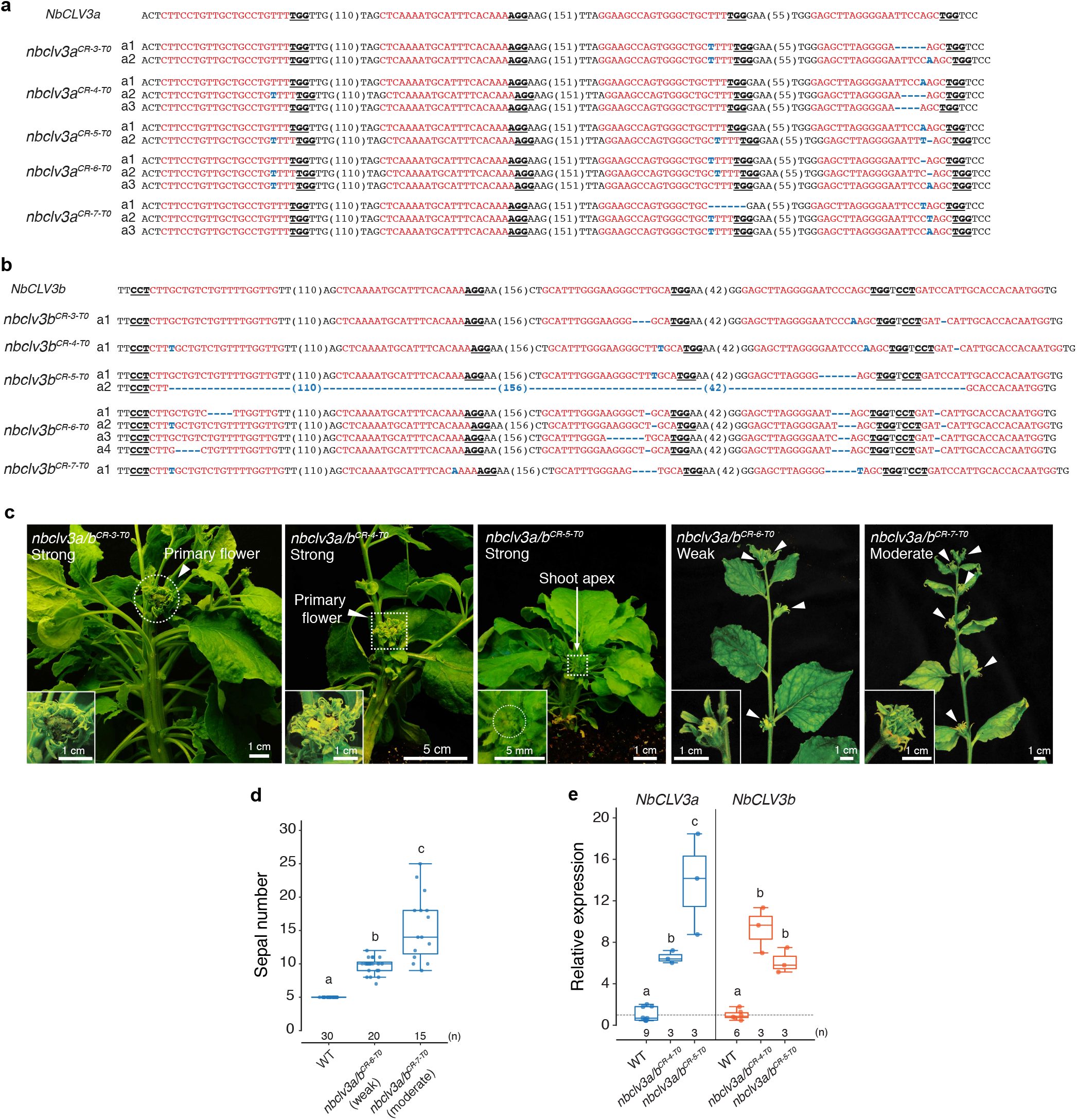
CRISPR-generated mutations of the tobacco *NbCLV3a* and *NbCLV3b* genes. **a**, CRISPR-generated sequences of *nbclv3a* mutant alleles. **b**, CRISPR-generated sequences of *nbclv3b* mutant alleles. Guide RNA and PAM sequences are highlighted in red and bold underlined, respectively. Blue letters and dashes indicate insertions and deletions, respectively. Numbers in parentheses represent gap lengths. **c**, Shoots and inflorescences of *nbclv3a/b* T_0_ plants. Three strong lines (*nbclv3a/b^CR-3-T0^, nbclv3a/b^CR-4-T0^* and *nbclv3a/b^CR-5-T0^*) show similar phenotypes compared to null *nbclv3a/b* mutants in **Fig. 1g**. Weak (*nbclv3a/b^CR-6-T0^*) and moderate (*nbclv3a/b^CR-7-T0^*) lines show regular shoot architecture but fasciated floral organs. White arrowheads indicate flowers. **d**, Sepal number of weak and moderate *nbclv3a/b* T_0_ plants. **e**, Relative expressions of *NbCLV3a* and *NbCLV3b* in shoot apices from *nbclv3a/b* T_0_ chimeric mutants, normalized to *NbPP2A*. Dashed line, value ‘1’ on the y-axis. Three and one biological replicates for WT and *nbclv3a/b* plants, respectively; two or three technical replicates included. Box plots, 25^th^-75^th^ percentile; center line, median; whiskers, full data range in **d** and **e**. The letters indicate the significance groups at *P* < 0.01 (One-way ANOVA and Tukey test) in **d** and **e**. The exact sample sizes (n) are shown as discrete numbers in **d** and **e**.

**Supplementary Fig. 2.**
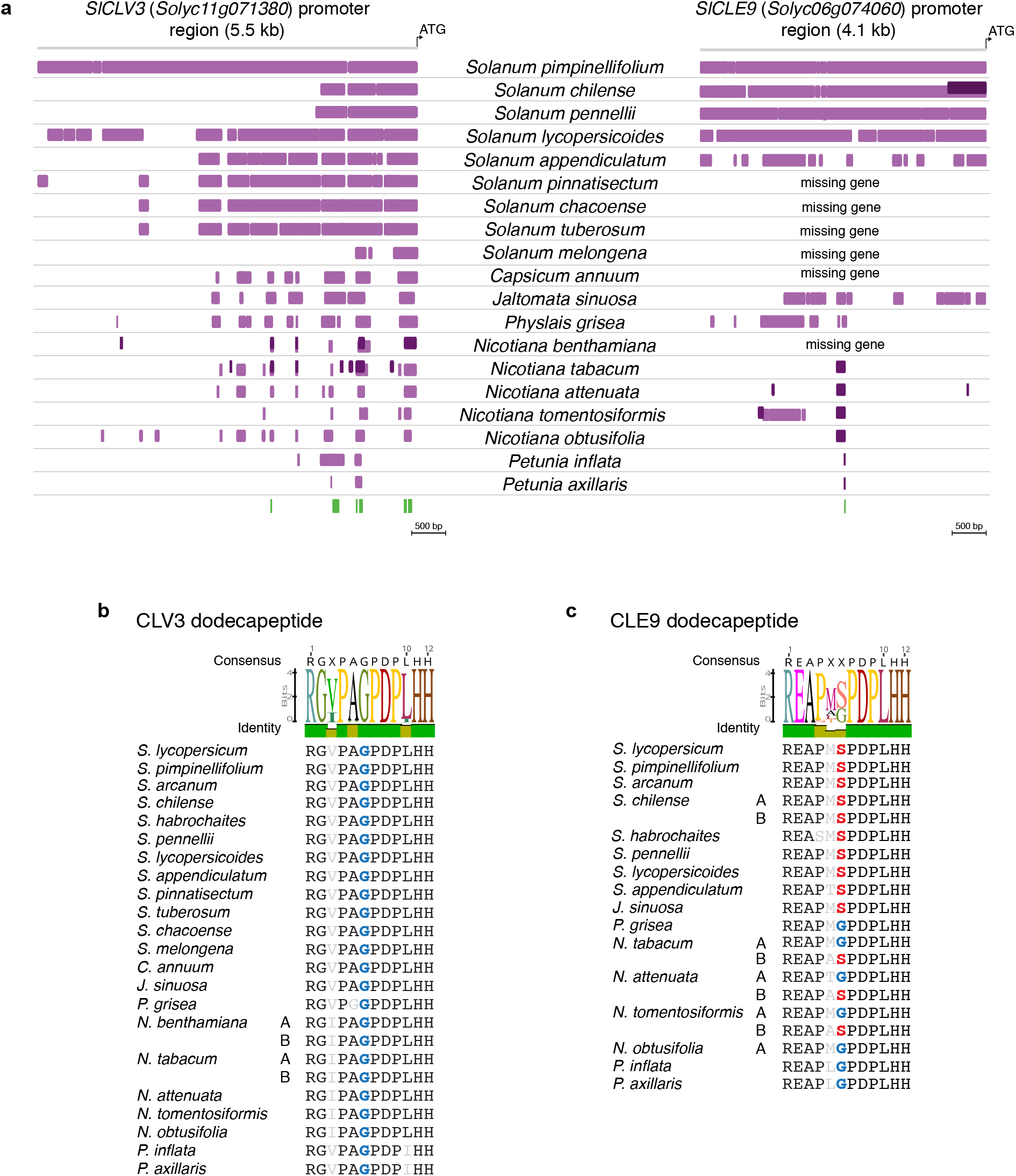
Conserved noncoding sequence (CNS) analysis of the promoter regions of *SlCLV3* and *SlCLE9* orthologs in the Solanaceae family. **a**, Conservatory analysis of Solanaceae *CLV3* and *CLE9* promoters. Purple boxes define highly similar regions of each gene’s orthologs in the indicated species, and dark purple boxes define highly similar regions of the paralogous gene (e.g. *CLV3B* or *CLE9B*) in the indicated species. Green boxes define Solanaceae CNSs. **b**, Multiple alignment and logo sequences of SlCLV3 dodecapeptide orthologs in the Solanaceae family. **c**, Multiple alignment and logo sequences of SlCLE9 dodecapeptide orthologs in the Solanaceae family.

**Supplementary Fig. 3.**
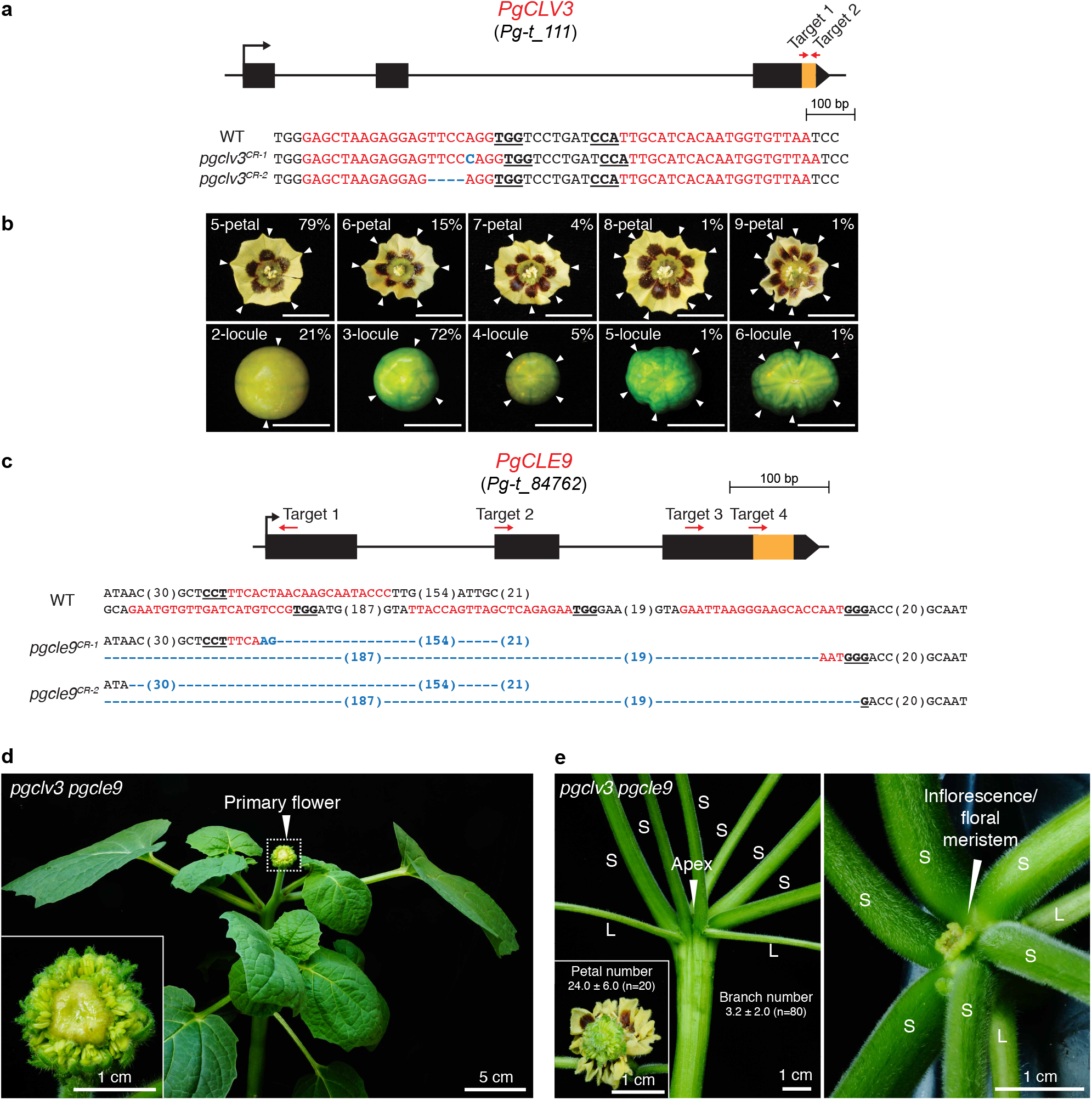
CRISPR-generated mutations of groundcherry *PgCLV3* and *PgCLE9*. **a**, Gene structure and sequences of *pgclv3* CRISPR mutants. **b**, Flowers and fruits of *pgclv3*. White arrowheads mark petals or locules. Percentages indicate the proportions of flower and fruit phenotypes. Scale bar, 1 cm. **c**, Gene structure and sequences of *pgcle9* CRISPR mutants. The orange rectangles in the gene structures indicate the regions of the CLE dodecapeptides in **a** and **c**. Guide RNA and PAM sequences are highlighted in red and bold underlined, respectively, in **a** and **c**. Blue letters and dashes indicate insertions and deletions, respectively, in **a** and **c**. Numbers in parentheses represent gap lengths in **a** and **c**. **d**, Shoot and an extremely fasciated primary flower of the *pgclv3 pgcle9* double mutant. **e**, Development of extra shoots (S) from the primary shoot and apex of a *pgclv3 pgcle9* double mutant. L, leaf petioles.

**Supplementary Fig. 4.**
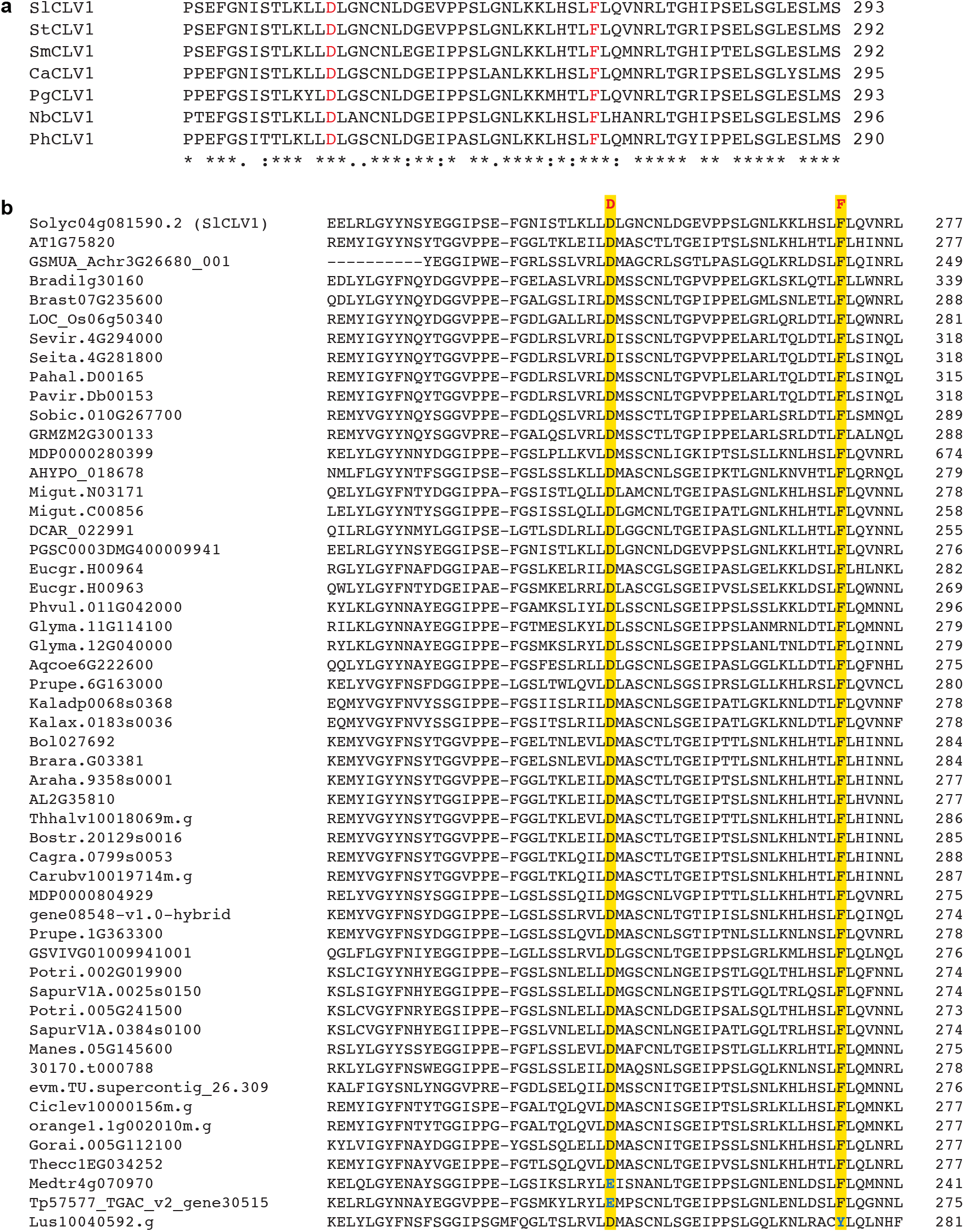
Sequence alignments of CLV1 receptor homologs. **a**, Alignment of the Solanaceae CLV1 protein sequences. Red letters indicate the two ultraconserved amino acids involved in the physical binding of CLE dodecapeptides. **b**, Alignment of CLV1 homologs in angiosperms. All the sequences are from the Phytozome v12.1 database (phytozome.jgi.doe.gov). Yellow highlights mark the conserved Asp and Phe. Detailed sequence information is shown in **Supplementary Dataset 3**.

**Supplementary Fig. 5.**
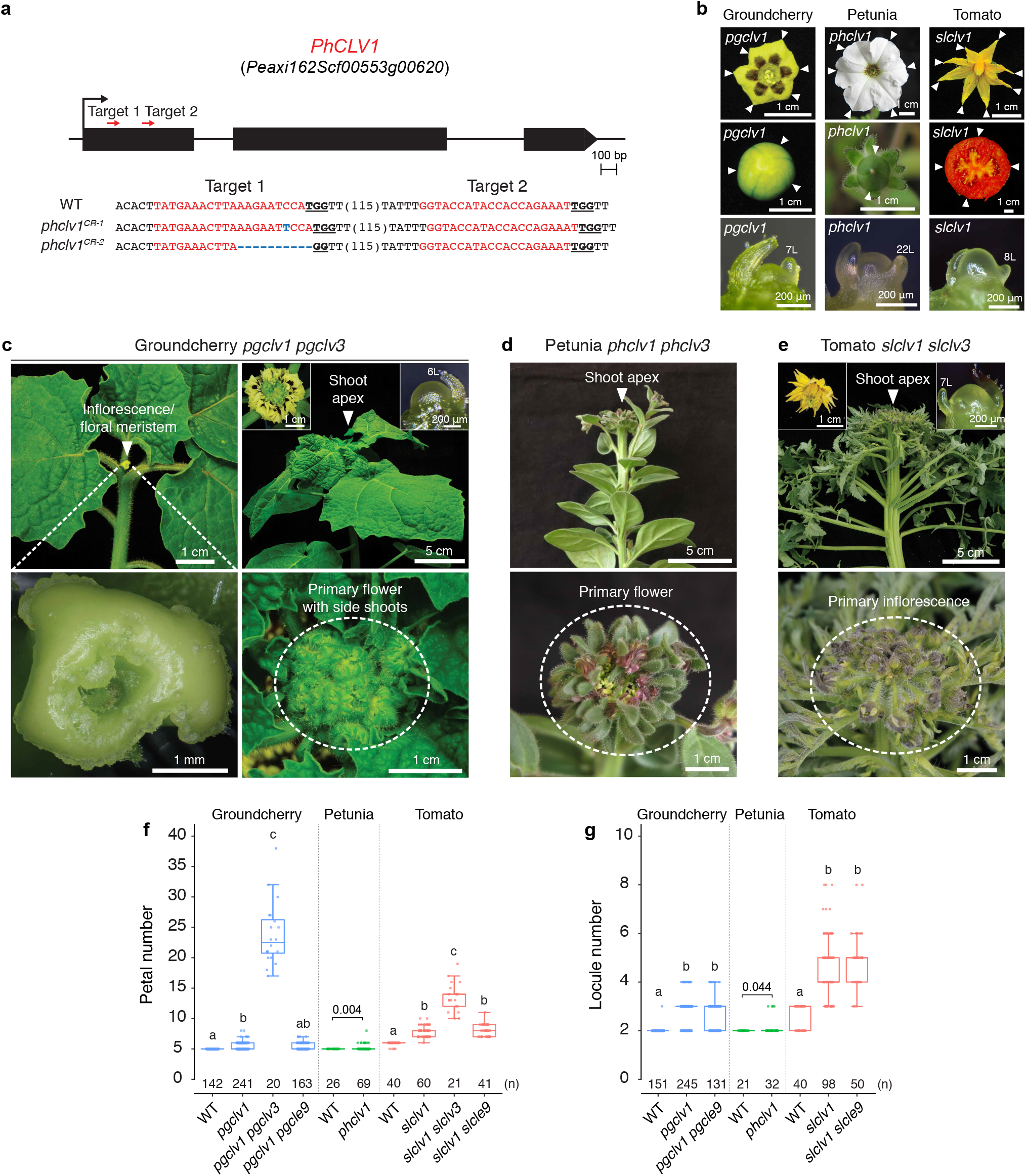
Groundcherry *pgclv1 pgclv3* and petunia *phclv1 phclv3* double mutants are severely fasciated like tomato *slclv1 slclv3* double mutants. **a**, Gene structure and sequences of two *phclv1* CRISPR mutants. Guide RNA and PAM sequences are highlighted in red and bold underlined, respectively. Blue letters and dashes indicate insertions and deletions, respectively. Numbers in parentheses represent gap lengths. **b**, Flowers, fruits/pods, and meristems from *pgclv1*, *phclv1*, and *slclv1* single mutants. White arrowheads mark petals or locules. 7L, 7^th^ leaf primordium, 8L, 8^th^ leaf primordium. 22L, 22^th^ leaf primordium. **C**, Side and top-down views of a *pgclv1 pgclv3* double mutant shoot, inflorescence/floral meristem, and primary inflorescence. 6L, 6^th^ leaf primordium. **D**, Side and top-down views of a *phclv1 phclv3* double mutant shoot and primary flower. **E**, Side and top-down views of a *slclv1 slclv3* double mutant shoot, flower, vegetative meristem and primary inflorescence. Fasciated flowers and vegetative meristems are shown in insets of **c** and **e**. **f**, **g**, Petal (**f**) and locule (**g**) numbers of groundcherry WT, *pgclv1, pgclv1 pgclv3, pgclv1 pgcle9*, and petunia WT, *phclv1*, and tomato WT, *slclv1, slclv1 slclv3*, and *slclv1 slcle9*. Not that all three Solanaceae *clv1* single mutant fasciation phenotypes are similarly weak. Box plots, 25^th^-75^th^ percentile; center line, median; whiskers, full data range in **d** and **e**. The letters indicate the significance groups at *P* < 0.01 (One-way ANOVA and Tukey test) in **f** and **g**. *P* values (two-tailed, two-sample *t*-test) in **f** and **g**. Exact sample sizes (n) are shown in **f** and **g**.

**Supplementary Fig. 6.**
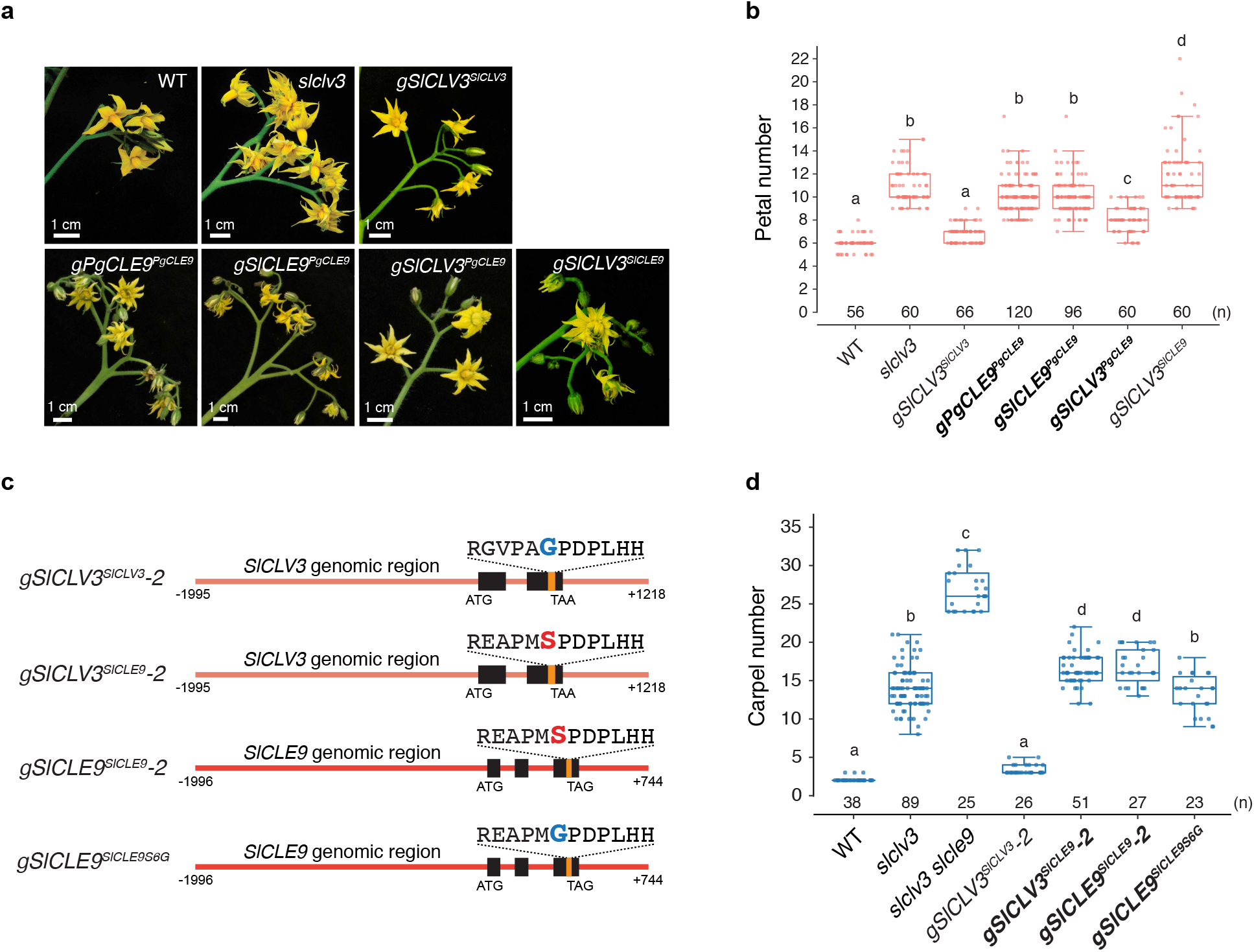
Transgenic complementation tests of tomato *slclv3* single and *slclv3 slcle9* double mutants. **a**, **b**, Complementation tests of tomato *slclv3* single mutants. Inflorescence images (**a**) and petal number quantifications (**b**) of WT and *slclv3* compared to the T_1_ transgenic plants *gSlCLV3^SlCLV3^*, *gPgCLE9^PgCLE9^, gSlCLE9^PgCLE9^, gSlCLV3^PgCLE9^*, and *gSlCLV3^SlCLE9^*. **c**, Diagrams of the constructs used for complementation tests of *slclv3 slcle9* double mutants. *gSlCLV3^SlCLV3^* (*SlCLV3* genomic DNA). *gSlCLV3^SlCLE9^* (*SlCLV3* genomic DNA including the sequence for SlCLE9 dodecapeptide). *gSlCLE9^SlCLE9^* (*SlCLE9* genomic DNA). *gSlCLE9^SlCLE9S6G^* (*SlCLE9* genomic DNA including the sequence for PgCLE9 dodecapeptide). Black and orange rectangles mark the coding sequences and the dodecapeptide sequences, respectively. The numbers with minus (-) and plus (+) signs indicate the positions of the upstream sequences and the downstream sequences from the adenines of start codons, respectively. **d**, Carpel number quantifications of WT, *slclv3, slclv3 slcle9* mutants compared to the T_1_ transgenic plants *gSlCLV3^SlCLV3^-2*, *gSlCLV3^SlCLE9^-2*, *gSlCLE9^SlCLE9^-2*, and *gSlCLE9^SlCLE9S6G^*. Data are based on at least three independent transgenic lines for each construct. Box plots, 25^th^-75^th^ percentile; center line, median; whiskers, full data range in **b** and **d**. The letters indicate the significance groups at *P* < 0.01 (One-way ANOVA and Tukey test) in **b** and **d**. Exact sample sizes (n) are shown in **b** and **d**.

### Supplementary Tables

**Supplementary Table 1.** CLE dodecapeptide sequences of SlCLV3 and SlCLE9 homologs

**Supplementary Table 2.** Differentially expressed genes between petunia WT and *phclv3* from mRNA-seq.

**Supplementary Table 3.** Differentially expressed genes between groundcherry WT and *pgclv3* from mRNA-seq.

**Supplementary Table 4.** Primers used in this study.

### Source data

**Source data 1.** CRISPR-generated mutations in this study

**Source data 2.** Raw data of relative expression for Supplementary Fig.1.

**Source data 3.** Quantification data for organ numbers in this study.

**Source data 4.** Quantification data for meristem size from Fig. 1, 2 and 3.

**Source data 5.** Normalized counts from mRNA-seq for Fig.2 and 3.

**Source data 6.** Syntenic region of *SlCLV3* homologs, defined by Conservatory orthogroups

**Source data 7.** Syntenic region of *SlCLE9* homologs, defined by Conservatory orthogroups

**Source data 8.** Exact *P*-values in this study (one-way ANOVA with Tukey test)

### Supplementary Dataset

**Supplementary Dataset 1.** Sequencing trace files.

**Supplementary Dataset 2.** Petunia PhCLE9 sequence.

**Supplementary Dataset 3.** CLV1 homolog sequences.

## Notes

### Competing Interest Statement

The authors have declared no competing interest.

